# Rigor and Transparency in two neurotrauma-publishing journals: editorial policies improve transparent reporting

**DOI:** 10.64898/2026.01.16.699952

**Authors:** Anita Bandrowski, Amit Namburi, Adam Ferguson, Candace L. Floyd, Maryann E. Martone

## Abstract

Preclinical research in traumatic brain injury (TBI) continues to significantly increase knowledge and yield a large number of peer-reviewed studies, but translation of these results to the clinical setting has been minimal. Rigor and transparency factors such as concealment of group allocation (e.g., “blinding’’) or ensuring that reagents are identifiable are critical in ensuring that scientific studies are replicable and translatable. Yet, nearly all efforts aimed at measuring these factors have concluded that reporting practices are problematic and incomplete. One way to improve transparency of reporting practices is to require that authors address a set of transparency related items in some way, such as a checklist or a paper section. Recently, *Journal of Neurotrauma*, a leading publisher of preclinical TBI research, instituted a required rigor-related section, which is explained to authors via a set of transparency, rigor, and reproducibility (TRR) instructions (one example for each manuscript type). These documents include specific transparency sections explaining blinding, power calculations, protocols, code, and data deposition. *Experimental Neurology* is a journal that is similar in size, impact and topic but the journal does not have explicit instructions to authors about transparency items. The purpose of this study was to assess the degree to which transparency reporting items were included in published manuscripts comparing reporting practices in the *Journal of Neurotrauma* and *Experimental Neurology*. We used a commercial software, SciScore, which is an AI tool tuned to detect rigor/transparency sentences in published manuscripts and count the number found (roughly dividing by the number expected) to obtain a score. Overall, SciScore found that in 6 of 8 items that were explicitly asked for, such as power calculations, investigator blinding, inclusion criteria, attrition, and data were significantly greater (more than 10%) compared to *Experimental Neurology*. However in *Journal of Neurotrauma* papers with the extra rigor section, 3 of 4 rigor items that were not explicitly asked for in the template rigor documents, such as subject demographics or transparent antibody reporting were not different from *Experimental Neurology*. One item, reporting of the sex of subjects was significantly better in *Experimental Neurology*. This shows that the *Journal of Neurotrauma* required rigor section is effective in improving reporting, but it would be far better if sex as a biological variable and transparent reporting of reagents (items present on major checklists including NIH rigor criteria) would be included.

## Introduction

Traumatic Brain Injury (TBI) is a major cause of death and long term disability, impacting nearly 50 million individuals per year, and costing the global economy $400B/year (Maas et al., 2022). Unfortunately, as of 2025 all randomized controlled trials of TBI therapeutics have failed to demonstrate efficacy in their primary endpoint (Lynch et al., 2023; Guo et al., 2023), despite initial promise in laboratory (preclinical) models. This failure of translation from bench to bedside is complex and can be attributed to a myriad of reasons, however the lack of standardization and reproducibility in laboratory studies of TBI likely contributes to these translational challenges.

Transparent reporting of experimental conditions and rigorous practices have been highlighted by the National Institutes of Health and other major organizations as key sources of irreproducibility of studies (Landis et al, 2012). A plan for addressing four of these practices has been required with each grant submission since 2016 (NIH NOT-OD-15-103): the lack of randomization, the lack of investigator blinding, the lack of verification of group size (power calculations), and the lack of authentication of key biological and chemical resources such as antibodies, cell lines, and transgenic organisms. Other factors including the full description of inclusion/exclusion criteria, attrition of subjects, biological variables like sex, and statistical issues are also emphasized as being good practice for preclinical studies. While ensuring that a study is blinded or properly randomized does not guarantee that it will be reproducible, there is very good evidence that randomization and blinding reduce the effect size of any treatment by about half (Macleod et al, 2004; 2015). Therefore studies that are marginally significant without blinding will likely not be significant if investigators were naive to the treatment condition. The PRECISE TBI (PRE Clinical Interagency reSearch resourcE-Traumatic Brain Injury, http://precise-tbi.org, RRID:SCR_022252) group has been charged with aiding the preclinical TBI community to implement these standards and to submit data to a dedicated repository ODC-TBI.org (RRID:SCR_021736). These efforts include encouraging the community towards better reporting practices, however we have not assessed, until now, whether the community is already following these practices.

Various scientific societies and research journals (Nature Checklist 2018, Marcus et al, 2016; MacLeod 2015, McNutt et al, 2014), including the *Journal of Neurotrauma*, have adopted guidelines to improve rigor and reproducibility practices in their journals. Since 2022 the *Journal of Neurotrauma* specifically asks authors to include TRR information in a separate section dedicated to rigorous reporting practices (Editorial Policy, *Journal of Neurotrauma*). The journal considers this section required. This is akin to asking authors to include a “Data Availability” section, which is a common practice in many journals, but as of the writing of this manuscript there was no direct study pointing to a specific quantifiable impact on author compliance of the presence of this section (many attempts have been made to quantify data sharing, see Woods & Pinfield, 2022; Baxter et al, 2024; Parkin et al, 2019; Riedel et al, 2020). On the other hand, other journals, e.g., *Experimental Neurology*, have not taken those steps. Importantly, it is incorrect to assume that checklists and editorial policies directly lead to a change in practice for conducting research in a rigorous manner or even reporting research in a transparent manner. For example, authors who acknowledged the ARRIVE guidelines (Animals in Research: Reporting In Vivo Experiments, National Center for the Replacement, Refinement and Reduction of Animals in Research, 2020) in their manuscripts, as required by PLoS One journal policy, did not actually report more ARRIVE transparency items than their counterparts who ignored the guideline (Leung et al, 2018). Therefore, it is important to further understand the true impact of guidelines on publication practices.

In this study we looked at two journals that publish reports from the same community of researchers: The *Journal of Neurotrauma* (JNeuroT) and *Experimental Neurology* (ExpNeuro). These journals were selected for comparison due to similar size in terms of the number of papers published annually, coverage of many of the same topics, and similarities in journal impact factor numbers (2024 according to SciMago, RRID:SCR_027328). There are differences in their editorial practices, especially recently with the advent of the TRR section, implemented in 2022 in JNeuroT. These differences in editorial practices set up a natural experiment that has three groups: JNeuroT prior to the TTR statement requirement, JNeuroT with the extra rigor section (JNeuroT+) and ExpNeuro as a control.

This study employed an innovative approach. We used the AI-based SciScore tool to evaluate the adherence of all evaluated studies to pre-established rigor criteria. This enabled us to overcome limitations of other approaches which often rely on selection of a reasonably small, representative sample for manual evaluation, which is highly labor intensive. Our use of the AI based SciScore tool facilitated our evaluation of all published papers in the time period under consideration and compared these to a known false positive and false negative error rate (Menke et al., 2020, 2022), eliminating the need to sample papers from other journals.

This study analyzed full-text research articles published in *Experimental Neurology (ExpNeuro)* and the *Journal of Neurotrauma (JNeuroT without extra section or JNeuroT+ with extra section)* between 2014 and 2024, filtered for the subset present in PubMed Central (PMC; see supplementary file). Articles were included if they had a valid PubMed ID (PMID) that could be successfully converted to a PubMed Central ID (PMCID), enabling access to the full article text for automated analysis.

## Methods

The method section for each manuscript and the additional rigor section, when applicable, were submitted to SciScore (Version 2, RRID:SCR_016251), which parses the text provided and matches each sentence and part of sentence to one or more rigor criteria including randomization, blinding, power calculations, presence of data, presence of code, and the presence and findability of each key biological resource, with F1 scores, the harmonic mean of precision and recall, and a more broadly defined accuracy rate reported previously (Menke et al 2022). SciScore assigns a score of 1-10, reflecting adherence to rigor criteria (see Table 1). For nearly all analyses, we exclude manuscripts that had no methods section or those that scored a 0 so that we can compare the overall scores with the Rigor and Transparency Index, reported by Menke, 2022. Menke and colleagues calculated scores for over 2 million papers in the open access subset of PubMed Central. In the Menke paper, any papers that scored a 0 were excluded. Zero scoring papers may be review papers without a methods section, they may be papers in which the methods section is included in the supplement, or they may be papers in which no rigor criterion or reagent is present. For the current study, we are using the SciScores for the 331,393 papers from the year 2020 because this is the most recent year thus most comparable to the current analysis.

**Table 1:**
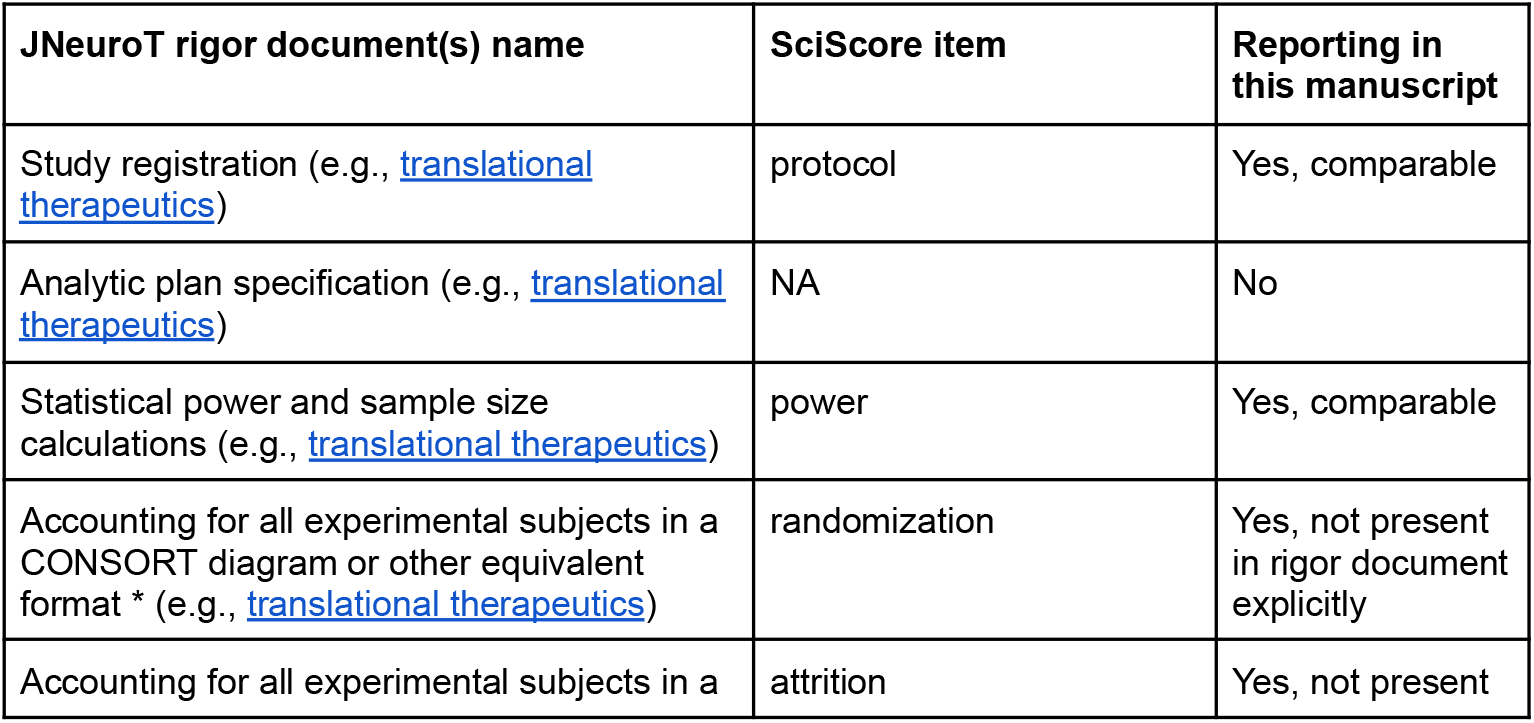
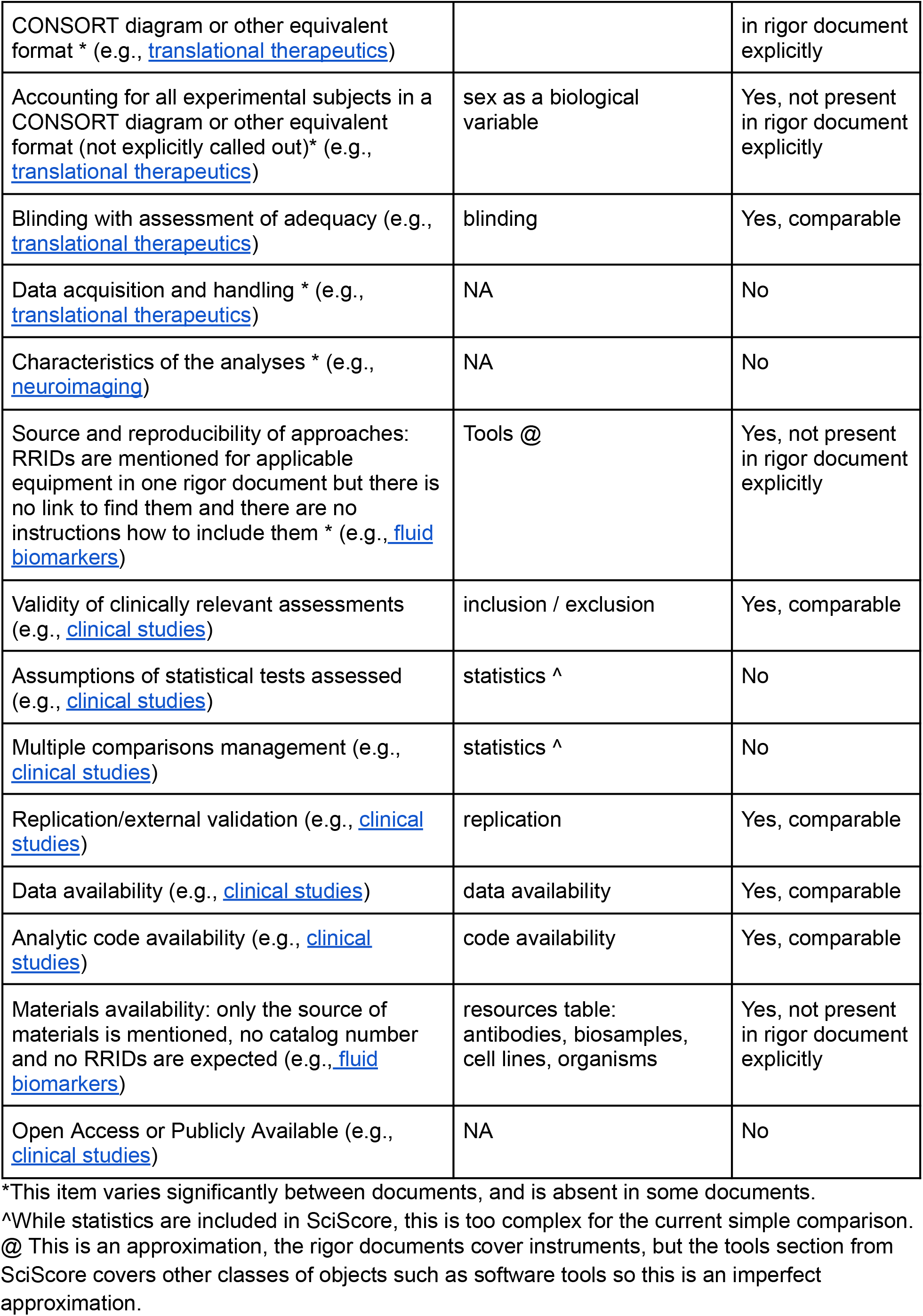
A comparison of the JNeuroT rigor document items and the rigor items verified by the SciScore tool. The third column notes whether the item is reported in the current study. Examples lead to one example from one of the TRR documents but please note, these items are usually present in many TRR documents accessible on the Journal of Neurotrauma author instructions “preparing your manuscript” section.

### Identifier Mapping

To obtain PMCIDs corresponding to each PMID, we used the NCBI ID Converter API RRID:SCR_013249, https://www.ncbi.nlm.nih.gov/pmc/utils/idconv/v1.0/). This service supports programmatic resolution between article identifiers and was accessed in batch mode to convert the full list of PMIDs to their associated PMCIDs. Articles without a matching PMCID or without full-text availability in PMC were excluded from further analysis.

### Full-Text Retrieval

For each article with a valid PMCID, full-text HTML content was retrieved from the PMC website. Articles were accessed using the canonical PMC URL structure (e.g., https://pmc.ncbi.nlm.nih.gov/articles/PMC11265769/). The retrieval process was implemented in Python RRID:SCR_008394, using the requests library RRID:SCR_026988 for web access and BeautifulSoup4 (RRID:SCR_026987) for HTML parsing.

### Section Extraction

Two types of content were extracted from each article (all papers, regardless of type, were extracted):

#### Methods Section

For all included articles, we extracted the section describing the study’s experimental procedures. Candidate sections were identified by searching for headings containing common variants such as “Methods,” “Materials and Methods,” or “Methodology.” Once identified, the accompanying body text was parsed and stored (see supplement).

#### Transparency, Rigor and Reproducibility Section

In the *Journal of Neurotrauma*, articles submitted on or after January 1, 2023, were expected to include an additional rigor section. Therefore, we searched for a dedicated section on transparency, rigor, and reproducibility or TRR. Headings were identified using a keyword-based approach, and the corresponding text was extracted in full when present (see supplemental file for the extracted text - Extra_Sections.csv).

### Data Organization and Output

All extracted content was saved in JSON format (see supplement). Each record included the PMCID, publication year (parsed article metadata), and the extracted Methods and, when applicable, Rigor sections. Articles for which extraction failed or relevant sections could not be found were logged separately for transparency and error tracking.

### Comparison of SciScore and the JNeuroT Rigor Documents

We examined the 7 rigor documents listed in the instructions to authors at JNeuroT. Items that are also detected by SciScore are listed in Table 1. While the rigor documents specify two separate questions dealing with study materials, it should be noted that the example sentences provided would not be considered sufficient by SciScore because SciScore was tuned to detect specific resources such as a particular software tool like ImageJ, a particular instrument or a particular mouse, and whether or not that item can be found in the relevant catalog. Statements such as those proposed by the rigor documents such as “*All materials used to conduct the study were obtained from a widely available source:_________(provide reference)*.” (direct link in Table 1) would be considered as not fulfilling the requirement to list the reagents used in a manner that is findable as in FAIR, as checked by SciScore. Therefore, the guidelines for JNeuroT and SciScore agree in principle but not in the details. SciScore also determines if the item is not detected but expected (“not detected”) or not detected but not expected (“not required”). For the current analysis we only examined the items that are expected but not found.

### Statistics

We evaluated statistical significance using a two-sample Z-test for two population proportions using the assumption that p < 0.05 (two-tailed) using the Social Science Statistics Calculator, RRID:SCR_016762. We tested the differences between JNeuroT+ vs JNeuroT, and JNeuroT+ vs ExpNeurol (Results in Table 3). We added the numbers for each criterion from the Menke (2022) dataset as a benchmark comparison, but we did not seek to determine if these were significantly different. Each Z score, exact p value is reported, with red text indicating not significant at p > 0.05, and blue for p < 0.05.

## Code and Data Availability

All scripts (code written specifically for this task) used for identifier conversion, full-text retrieval, and section parsing are publicly available in a dedicated repository https://github.com/namburiamit/pubmed-section-extraction DOI:10.5281/zenodo.15430521

- GitHub link for the extracted data and its code: https://github.com/namburiamit/pubmed-section-extraction/tree/main/Data/Extracted%20Experiment%20Set
- Data in JSON files for Experimental Neurology: https://github.com/namburiamit/pubmed-section-extraction/tree/main/Data/Extracted%20Jneurotrauma-new
- Data (methods section) in JSON files for JNeuroT: https://github.com/namburiamit/pubmed-section-extraction/tree/main/Data/Additional%20Section%20-%20extracted

Additional Section (Transparency, Rigor, and Reproducibility Statement) in JSON files for JNeuroT Spreadsheet of the data for both of the journals: https://docs.google.com/spreadsheets/d/1DQUG4GE5H3STMC6JTRS1vRZseLj3agjGHQ0sw6BRXgs/edit?usp=sharing

Data analysis for this manuscript is included here: https://docs.google.com/spreadsheets/d/1DQUG4GE5H3STMC6JTRS1vRZseLj3agjGHQ0sw6BRXgs/edit?usp=sharing

## Results

Our analysis of all papers published regardless of type and deposited to PubMed Central in ExpNeurol and JNeuroT (publication years 2014-2024) showed a few important trends. First, as the JNeuroT policy on reproducibility documentation went into effect on January 1 2023 (Editorial Policy *Journal of Neurotrauma*) for all submissions, we expected that papers in 2023, and 2024 should increasingly have the “extra section” as per instructions to authors, based on an average time to initial decision of 25 days, with processing of manuscripts also taking some time. Our dataset shows that the JNeuroT papers published in 2023 included 43 manuscripts without the extra section and 20 with extra section (JNeuroT+) and in 2024 only one manuscript had no extra section and 8 manuscripts (JNeuroT+) had the extra section. The 2024 data is partial because it was both extracted in 2024, and manuscripts take time to be added to PubMedCentral, making the most recent year, also the most incomplete. We also note that several papers from years before 2023 had a rigor section included.

Figure 1 shows that JNeuroT is slightly higher scoring than ExpNeuro in all but 3 years. The JNeuroT+ scores are highest, but in 2022 while the absolute number is most different, the significance of this number is questionable because there were only two papers with the extra section, making any conclusion suspect. In 2023, where a reasonably high 23 papers with extra section vs 43 papers without extra section is likely to be meaningful. The 2024 data are also unlikely to be meaningful because there is only a single manuscript without an extra section. Interestingly, the score for ExpNeuro is exactly the same as JNeuroT, a rare occurrence. We are not aware if there are policy changes at the journal that might have driven this change.

**Figure 1:**
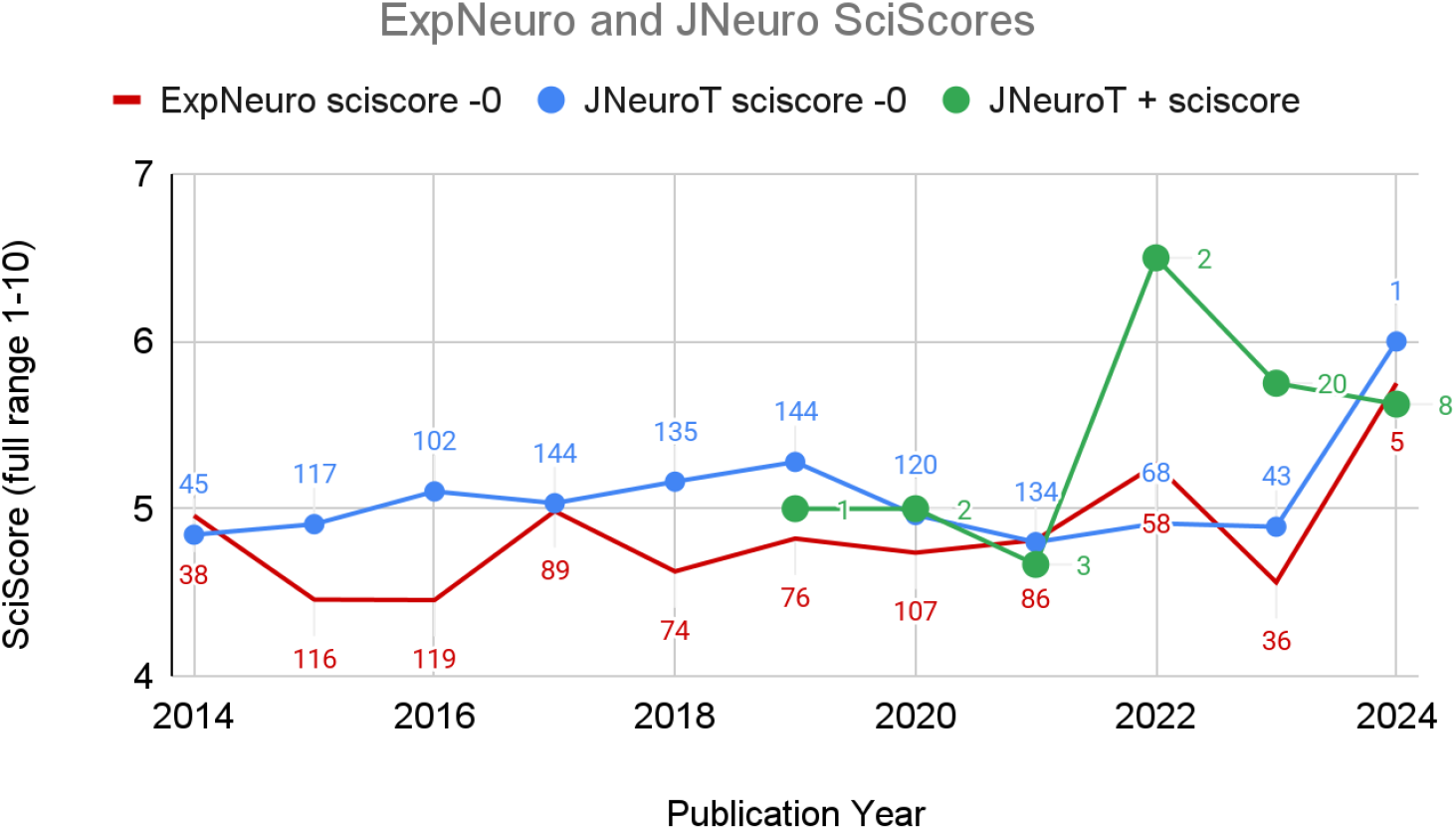
Chart of papers analyzed using SciScore over time. The “-0” indicates that papers scoring 0 were not included in the analysis. The numbers next to each point represent the number of papers that were analyzed in the category (per Journal per Year).

The results of SciScore’s individual items showed multiple significant differences. ExpNeuro scores are generally lower than JNeuroT scores, and within JNeuroT, the scores of rigor section (JNeuroT+) papers are higher. All groups were also significantly different from the overall PubMed Central average scores for the 2020 data based on 331,393 papers (previously published in Menke et al, 2022). Those authors who publish in JNeuroT and comply with adding the rigor section are also more compliant with rigor items by nearly 1 point, roughly corresponding to the addressment of one additional rigor item. Figure 2 shows that rigor items that are explicitly addressed by the extra section (left side of the graph) show large increases in the percentage of papers describing them, for example power calculations and data availability statements are nearly 50% of JNeuroT+ papers (green bars) compared to ∼10% (also see Table 3 for exact values). Where rigor items are not explicitly addressed (right side of Figure 2) there is little difference between the JNeurT+ and the other groups.

**Figure 2:**
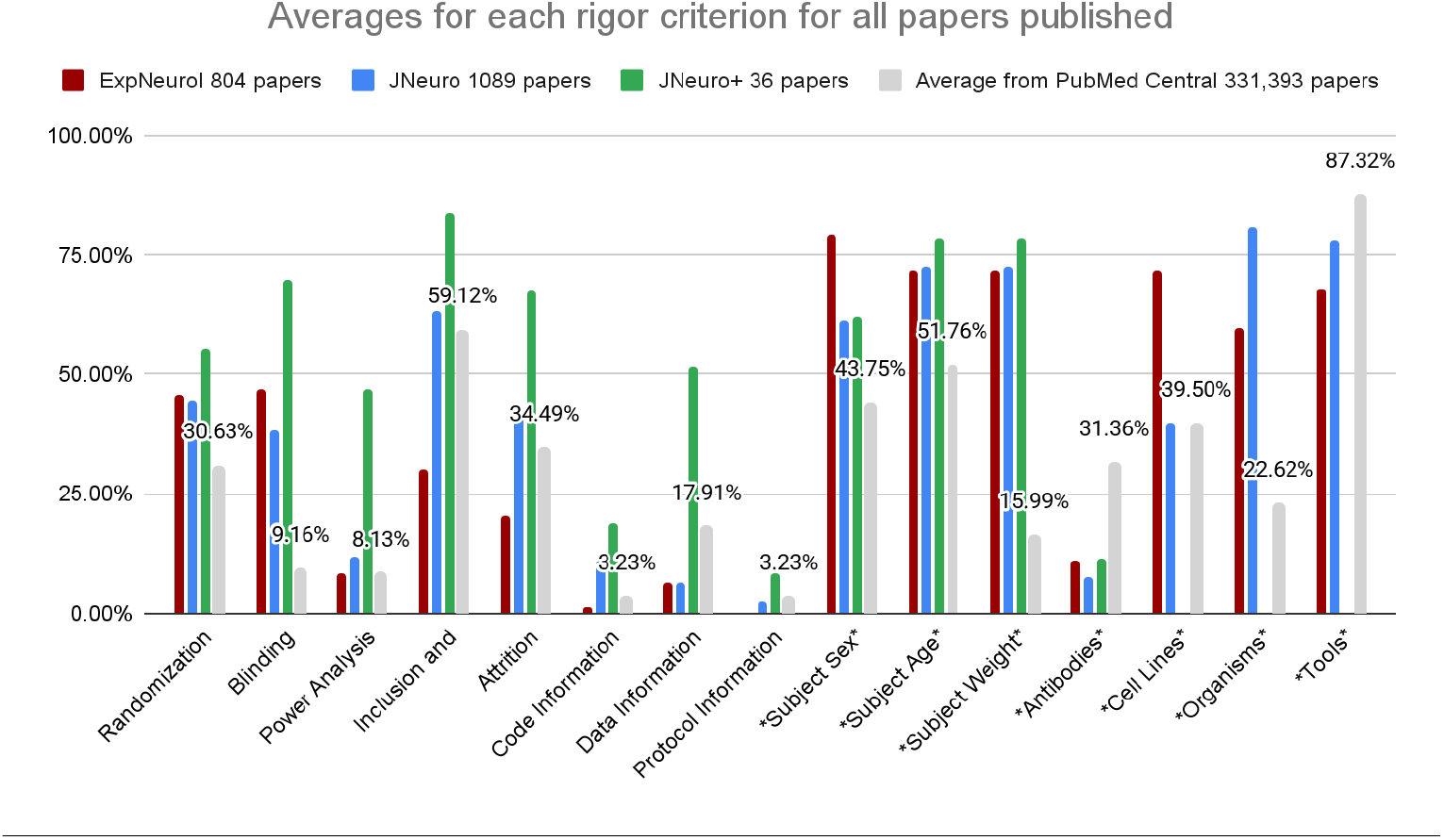
Chart of papers analyzed using SciScore, broken down by category. Labels with * are not present as a section on the JNeuroT+ rigor document; they may be mentioned as part of another criterion. Papers scoring 0 were not included in the analysis. The numbers next to each grouping represent the average of this criterion from 2020 papers as reported in Menke et al (2022, n = 331,393). Both journals are better than average in reporting randomization, blinding, subject demographics, and findable organisms, but they are worse than average when it comes to antibodies and tools. JNeuroT+ shows far higher percentages of reporting blinding, power analysis, attrition, and data, but antibody citations remain far lower than the average paper, while other resources such as organisms, cell lines and tools were not found in the 36 papers that also contained the extra rigor section.

Differences in overall scores between JNeuroT and ExpNeurol seem to be driven by author addressment of “inclusion and exclusion criteria” and attrition of subjects, as well as the inclusion of code. The drivers of JNeuroT+ (with rigor section) is higher for nearly all explicitly included criteria however the not explicitly addressed criteria such as sex, leads to reporting that is either no different or lower than ExpNeurol. Transparent reporting of antibodies is very low in both ExpNeurol and in JNeuroT+ around 10% while the average paper reports 30% of antibodies transparently (see Table 3). This suggests that rigor reporting of some criteria does not improve the manuscripts more generally, authors seem to largely stick to only what they are explicitly asked to address.

Identifiability of research resources such as antibodies, cell lines or organisms are included in major reproducibility documents such as the NIH rigor criteria, ARRIVE checklist for animal research and the MDAR checklist, but is not included in the JNeuroT rigor section. Unlike most top journals (Cell, Science, Nature, eLife) neither journal includes RRIDs, a major driving factor for findability of resources, in the instructions to authors (Bandrowski et al, 2015). The JNeuroT rigor document for Fluid Biomarkers does include one opaque reference to RRIDs but the other manuscript type sections do not. We consider these score drivers as “controls” and indeed, identifiability is lower for antibodies and tools compared to the 2020 PMC papers and roughly the same as ExpNeurol.

## Discussion

In our study, we found that rigor items are addressed by authors in the JNeuroT rigor section at a greater rate than the JNeuroT without rigor section or ExpNeurol, if the rigor items are in the rigor section documentation explicitly. It is possible that the presence of this additional section drives author behavior better than other ways of improving rigor, and this hypothesis is consistent with the Leung (2018) study looking at the effectiveness of the ARRIVE checklist in PLoS papers. Leung found that authors stating that they followed ARRIVE guidelines in PLoS papers were less effective than authors who published with the example JNeuroT+ TRR document (out of 27 ARRIVE items authors reported on average one additional item, so about a 3% improvement while the JNeuroT+ papers were routinely 10-20% better). Menke (2020) showed a larger change, a ∼20% improvement in the journal *Nature*, where the driver of the change was the *Nature* rigor checklist, which was enforced by editors. While authors do report more items when they are filling out a checklist, one might ask why do the checklists work so poorly overall? In the current experiment, one issue uncovered was that non explicit rigor items were reported at rates no different or worse than controls, so gains in one area cause losses in another. It may be that the rigor section gives authors a false sense of security that they are publishing rigorously and do not pay attention to other rigor items that they might normally pay attention to. Another potential issue is that we do not have sufficient data for whether the rigor documents lead to sustained improvements in reporting or whether they lead to a relatively small and non-durable spike in reporting. The 2024 data show that ExpNeuro overall scores are higher than the JNeuroT+, which is decreasing after an initial large improvement. It is unlikely that with small n’s for the 2022 timepoint the difference is significant, however the 2024 data, where the scores decrease from 2023 in JNeuroT+ are a troubling development. This will need to be followed up in several years to determine if the rigor documents are impactful over a longer term.

Rigor and reproducibility issues plague the scientific literature, especially the literature reporting on studies in TBI, stroke and spinal cord injury, which for many years produced no theories, practices or solutions that can underpin treatments or cures of these major disorders. Sharing of raw data (Fouad et al, 2020) led to one augmentation of treatment protocols for spinal cord injury that is showing promise in surgical settings. One would imagine that this type of result, a clinically significant finding based on the sharing of data, would drive the field and especially two major journals for preclinical TBI to mandate the inclusion of data in data repositories and other factors of transparency such as RRIDs in all journal articles as suggested by the PRECISE-TBI group (precise-tbi.org). Indeed both journals studied have higher rigor and transparency scores than the average journal, which likely reflects the topic (inclusion of many human studies, which are generally better than preclinical studies in terms of following rigor checklists, see Menke et al, 2020), but the difference is small, except in the 37 papers that follow the extra rigor guideline suggested by JNeuroT. These papers, within which authors followed the extra rigor document are far better, especially in the categories suggested by the document such as the deposition of data, calculating group size, and reporting on blinding of investigators or analysts.

### Limitations of this study

We acknowledge that although we have utilized a tool that can parse thousands of manuscripts, the number of manuscripts with the rigor document is still relatively small, 37. This is simply a factor of when we processed the data. We also must acknowledge that the study is limited to papers with additional rigor sections published over only about one year, which is limiting in that authors have not yet had a chance to learn from the rigor documents to implement the practices in their laboratories. For this reason, we did not attempt to further stratify the articles into bins such as “clinical” or “biomarkers” because this would make the statistics meaningless due to a very low number of papers of each type. The impact of the JNeuroT+ TRR documents may not yet be fully visible. We would also like to acknowledge that the software tool, SciScore has an accuracy rate that is less than 100% (above 90% in all cases other than subject demographics, for exact numbers for each category please see Table 2 Menke et al, 2022), therefore any one manuscript processed will have one or more false positives or false negatives. We did not seek to re-curate these specific manuscripts to determine if the previously reported rates of false positives and false negatives are accurate for this set of journals as that was out of scope.

**Table 2:** A summary of the journals examined. All raw data are included in the data analysis file. Zero scoring papers include types of papers that are not applicable such as reviews, but they may also include papers that should be scored but have no single rigor criterion mentioned. These are eliminated from subsequent analysis.

|  | ExpNeurol | JNeuroT | JNeuroT+ | PMC (2020) |
| --- | --- | --- | --- | --- |
| **Total papers** | 804 | 1089 | 37 | 331,393 |
| **SciScore** | 3.52 (SD 2.34) | 4.60 (SD 1.92) | 5.61 (SD 1.73) |  |
| **SciScore -zero** | 4.73 | 5.05 | 5.77 | 4.13 |
| **Number of zero scoring papers** | 206 | 96 | 1 |  |

**Table 3.** The results of analysis (data underlying Figure 2). Labels with ** and pink color are not present as a section on the JNeuroT+ rigor document; they may be mentioned as part of another criterion. Fields colored green represent rigor items that are explicitly listed on the rigor documents. Statistics reported here are comparing the proportion of the 37 papers in JNeuroT+ vs either the set of JNeuroT papers that do not have the rigor document or the full set of analyzed ExpNeurol papers. Significant results are blue, not significant results are red.

|  | ExpNeurol | JNeuroT | JNeuroT+ | PMC 2020 | JNeuroT+ vs JNeuroT | JNeuroT+ vs ExpNeurol |
| --- | --- | --- | --- | --- | --- | --- |
| **Total papers** | **804** | **1,089** | **37** | **331,393** |  |  |
| Randomization | 45.82% | 44.51% | 55.56% | 30.63% | z = -1.329.<br>p = .18352.<br>not significant at p < .05. | z = -1.1618.<br>p = .24604.<br>not significant at p < .05. |
| Blinding | 46.99% | 38.27% | 69.70% | 9.16% | z = -3.8494.<br>p = .00012.<br>significant at p < .05. | z = -2.6286.<br>p = .00854.<br>significant at p < .05. |
| Power Analysis | 8.36% | 11.78% | 47.06% | 8.13% | z = -6.2879.<br>p < .00001.<br>significant at p < .05. | z = -7.651.<br>p < .00001.<br>significant at p < .05. |
| Inclusion and Exclusion Criteria | 30.09% | 63.36% | 83.78% | 59.12% | z = -2.5453.<br>p = .01078.<br>significant at p < .05. | z = -6.8202.<br>p < .00001.<br>significant at p < .05. |
| Attrition | 20.30% | 41.03% | 67.57% | 34.49% | z = -3.2177.<br>p = .00128.<br>significant at p < .05. | z = -6.7453.<br>p < .00001.<br>significant at p < .05. |
| Code Information | 1.12% | 11.20% | 18.92% | 3.23% | z = -1.4939.<br>p = .13622.<br>not significant at p < .05. | z = -7.748.<br>p < .00001.<br>significant at p < .05. |
| Data Information | 6.22% | 6.43% | 51.35% | 17.91% | z = -9.9584.<br>p < .00001.<br>significant at p < .05. | z = -9.7799.<br>p < .00001.<br>significant at p < .05. |
| Protocol Information | 0.25% | 2.57% | 8.11% | 3.23% | z = -2.0258.<br>p = .04236.<br>significant at p < .05. | z = -2.0414.<br>p = .04136.<br>significant at p < .05. |
| *Subject Sex* | 79.07% | 61.35% | 62.16% | 43.75% | z = -0.0995.<br>p = .92034.<br>not significant at p < .05. | z = 2.4409.<br>p = .01468.<br>significant at p < .05. |
| *Subject Age* | 71.68% | 72.52% | 78.38% | 51.76% | z = -0.1153.<br>p = .90448.<br>not significant at p < .05. | z = -0.6595.<br>p = .50926.<br>not significant at p < .05. |
| *Subject Weight* | 71.68% | 72.52% | 78.38% | 15.99% | z = -0.1153.<br>p = .90448.<br>not significant at p < .05. | z = -0.6595.<br>p = .50926.<br>not significant at p < .05. |
| *Antibodies* | 11.19% | 7.57% | 11.42% | 31.36% | z = -0.9002.<br>p = .36812.<br>not significant at p < .05. | z = -0.0775.<br>p = .93624.<br>not significant at p < .05. |
| *Cell Lines* | 72.% | 40% | 0 | 39.50% |  |  |
| *Organisms* | 59.53% | 80.70% | 0 | 22.62% |  |  |
| *Tools* | 67.89% | 78.12% | 0 | 87.32% |  |  |

## Acknowledgements

We would like to thank the Veterans Administration for funding this work via VA I50BX005878. The contents do not represent the views of the U.S. Department of Veterans Affairs or the United States Government

